# Safety and efficacy of a new vaginal gel, Feminilove, for the treatment of symptoms associated with vaginal dryness and vulvovaginal atrophy in women: an *in vitro* and *in vivo* study

**DOI:** 10.1101/2021.08.17.455703

**Authors:** A Gonzalez, MR Lee, BA Johnson, L Booshehri, D Grady, V Vaddi, C Ip, C Mitchell, M Krychman, R Dardik

## Abstract

Vaginal dryness is a common symptom associated with vulvovaginal atrophy of menopause. The impact of vaginal dryness is very significant as it negatively affects quality of life, daily activities, sexual satisfaction as well as on interpersonal relationships. Symptoms of vaginal dryness is often underreported and undertreated. Recently, vaginal lubricants and moisturizers have been applied as one of the alternative and safe approaches to relieve vaginal dryness for women with mild to moderate vaginal dryness. We evaluated the safety and beneficial effects of a new type of estrogen-free vaginal gel, Feminilove BIO-FRESH moisturizing vaginal gel, using *in vitro* and *in vivo* experimental tools. Our results suggest that; 1) Feminilove vaginal gel exhibits minimal cell cytotoxicity on various human vaginal cells; 2) Feminilove vaginal gel exhibits minimal side-effects on the structure of vaginal mucosa stratum of experimental animals; 3) Feminiove vaginal gel inhibits the growth of pathogenic vaginal bacteria (*E. coli*) while promotes the growth of beneficial vaginal bacteria (*Lactobacillus spp)*; 4) Feminilove vaginal gel elicits an anti-inflammatory response on vaginal epithelial cells; and 5) Feminilove vaginal gel promotes the production of tropoelastin and collagen on cultural vaginal smooth muscle and may restore loose vaginal wall (*i.e.*, tightening effects). In summary, our results indicate that Feminilove BIO-FRESH moisturizing vaginal gel is a safe and effective remedy for vaginal dryness and vulvovaginal atrophy in women.

## I. INTRODUCTION

Vaginal dryness is common condition in women during and after menopause.^1–4^ Symptoms pf vaginal dryness has been associated with vaginal or vulvovaginal atrophy (VVA). ^5,6^ Prevalence of vaginal dryness due to VVA ranges from 15% to as high as 57% among postmenopausal women.^3^ A lack of natural vaginal lubrication during intercourse is an unpleasant experience for both sexual partners. As part of the cultural norms, men prefer and expect their female sexual partners to secrete vaginal lubrication during intercourse.^7^ In addition, sexually active women prefer to maintain and achieve proper vaginal lubrication during intercourse to enjoy the sexual activity as well as interpersonal intimacy.^8,9^ The underlying culprits for diminished vaginal lubrication may include hormonal changes due to menopause, stress, vaginal inflammation, chronic illnesses such as diabetes and chronic kidney disease, and medical treatment related side effects such as anti-depressants, chemotherapy and radiation.^9–11^ Vaginal dryness will likely lead to painful intercourse which has affected more than half of sexually active women at some point in their lives.^12–14^ Vaginal dryness and VVA has affected more than half of women during and after menopause.^15,16^ Recent survey in the US showed that vaginal dryness could also occur in younger women due to the utilization of antiestrogen medications.^17^ Symptoms of vaginal dryness and VVA are progressive and a variety of treatments include vaginal estrogen medication and different over the counter regimens such as vaginal moisturizers and lubricants.^18,19^ Recent data suggest that vaginal moisturizers and lubricants provide fast relief for vaginal dryness and dyspareunia and may attenuate discomfort for painful intercourse.^20–22^ Feminilove BIO-FRESH moisturizing vaginal gel, a new type of estrogen-free vaginal gel, was manufactured and distributed by Epoch NE Corporation, Seattle, WA, USA and Certified to NSF/ANSI 305 by Oregon Tilth. In this study, we aim to investigate the safety and efficacy of Feminilove vaginal gel for the treatment of symptoms associated with vaginal dryness and VVA by employing both *in vitro* and *in vivo* experimental models.

## II. METHODS

### 1. Cell viability test

We used the Cell Titer-Glo 2.0 Assay to assess cell viability.^23^ Actual procedures were according to the manufacturer’s guideline. We test the viability of 3 most popular vaginal gel products from Amazon (Vmagic Organic Vulva Cream vaginal moisturizer, Vulvacare vaginal moisturizer & intimate skin cream, Membrasin Topical Vulua Cream for Feminine Dryness) as well as Feminilove vaginal gel on 3 human cell lines (vaginal epithelial cell, cervical epithelial and endometrial epithelial cell). The detailed components for each product are listed in Table 1.

**Table 1:**
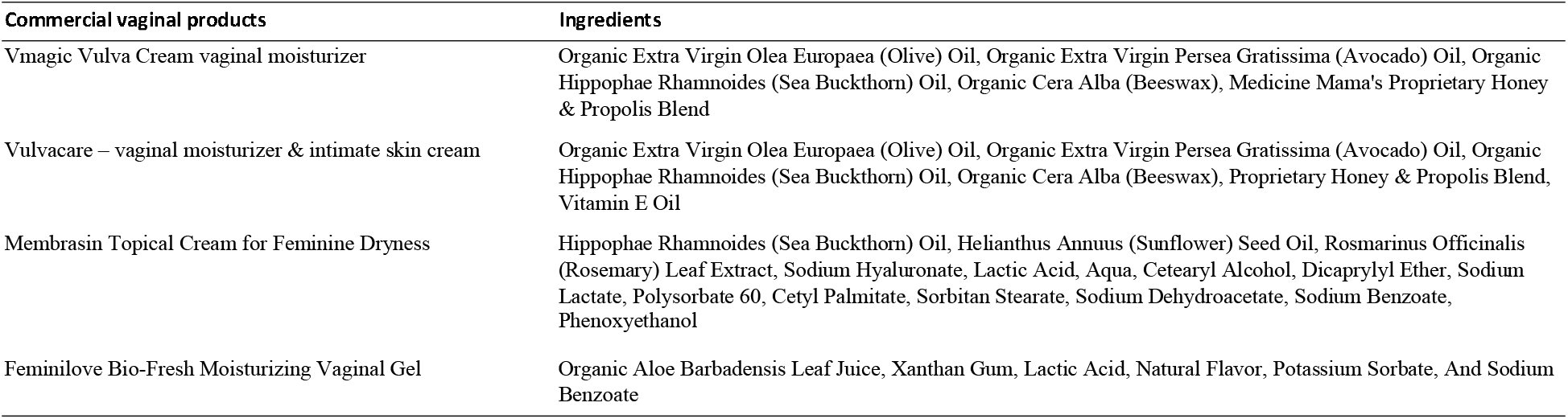
The detailed components for each vaginal moisturizer.

### 2. Animals

Sexually mature female New Zealand white rabbits (6-month-old) were purchased from Charles River Laboratories. All rabbits were fed *ad libitum* with rabbit food pellets (Teklad 7015; Harlan Teklad). The acclimation is 4 weeks prior to the use in the intravaginal study. Procedures listed in this study were strictly followed guidelines of USDA Animal Welfare Act and Animal Welfare Regulations.

#### 2.1 Rabbit vaginal irritation test

Female rabbits received 1 ml of placebo control gel or 1 ml of various Feminilove vaginal gel (0.01%, 0.1%, or 1.0%) via intravaginal administration for 10 consecutive days. The actual procedures for intravaginal instillation and insertion of the control gel or various contents of Feminilove vaginal gel were followed the published protocols.^24,25^

During the 10 days of study, we daily monitored the rabbits. On day 11, rabbits were sacrificed with injecting of ketamine and xylazine and serum from rabbits as well as vaginal tissues were extracted for subsequent investigation.

#### 2.2 Rabbit vaginal tissue histology evaluation

Fixed vaginal tissues were embedded and stained, following standard hematoxylin and eosin procedures. Slides of the vaginal sections were examined under microscope. Sections of vaginal tissues were evaluated for vaginal irritation for the scale of scores. 1 = no irritation; 2 = mild irritation; 3 = moderate and 4 = severe irritation.^24^ The sign of vaginal irritation includes exfoliation, infiltration of leukocyte and edema of vaginal epithelial cells.

#### 2.3 Cell proliferation

Rabbit vaginal sections were evaluated for cell proliferation (Zymed PCNA kit, Zymed Laboratories). Actual immunostaining Procedures were followed the manufacturer’s protocol. Stained vaginal sections observed under microscope and the scoring of a minimum of 600 cells was calculated.^24,25^

#### 2.4 Parameters of serum and blood chemistry

Blood samples from rabbits were analyzed, following standard procedrues.^25^ The hematology parameters include red blood cell count, leukocyte count, concentration of hemoglobin and hematocrit, platelet, and platelet volume.

### 3. *in vitro* reconstituted vaginal epithelium model

Three-dimensional tissue composed of normal human ectocervico-vaginal epithelial cells grown on polycarbonate filters using specially formulated serum-free medium were obtained from SkinEthic Laboratories (Nice, France). When cultured *in vitro* on a polycarbonate filter at the air-liquid interface in a defined medium, these cells form a 3-dimensional epithelial tissue resembling human *in vivo* vaginal mucosa. This *in vitro* tissue reproduces many of the histological, ultrastructural, and protein expression properties of native tissue, including interdigitation of cells, glycogen production, and cytokeratin expression. Before treatment, the tissue cultures were pre-incubated at 37 ^⍰^C and 5% CO_2_ for 24 hours. Prior to dosing, the medium was aspirated and triplicate wells of 6-well plates containing the *in vitro* reconstituted human vaginal epithelial tissues (size 0.5 cm^2^) grown on polycarbonate membrane filters were treated with or without increasing concentrations of FeminiLove (0.01, 0.1, or 1.0%) in 1 ml of medium or applied topically via a gel formulation and incubated for an additional 24 hours. The parameter determined *in vitro* was the histoarchitecture of exposed epithelial cells. At the end of the treatments, vaginal tissue inserts were washed in phosphate-buffered saline, and the polycarbonate filters covered with the vaginal tissue were cut out, fixed in 10% buffered formalin, embedded in paraffin, sectioned to 4 μm, and stained with hematoxylin and eosin as described previously.

### 4. Epithelial cell culture and stimulation

We purchased 3 human epithelial cells lines (vaginal cell line VK2/E6E7, endocervical End/E6E7 and ectocervical Ect/E6E7) from ATCC for this study. All cell lines were maintained in serum free keratinocyte medium and supplements with 0.5 mg/ml bovine serum, 0.1 ng/ml recombinant human epithelial growth factor, 50 mg/ml streptomycin and 50 U/ml penicillin (Life Technologies). Cultured cells were seeded into 24-well tissue culture plates for 7 days. Epithelial monolayers were co-cultured with 0.3% Feminilove in the presence or absence of the lipopolysaccharide (1 mg/ml) for 18 hours. prior to subsequent cytokine analysis. The production of IL-1β, IL-6, and TNFα from stimulated epithelial cells was assessed on a Bio-Plex 200 Luminex reader.

### 5. Fastin elastin assay

Human vaginal smooth muscle cells were grown to the confluence in a 96-well culture plates with 10% FBS. Culture medium were pre-heated to 65 °C prior to placing into the culture plates. We collected cell lysates and supernatants and cell lysates 48 hours after the initiation of experiment. Fastin Elastin Assay kits (Biocolor Ltd, Carrickfergus, UK was used to assess the contents of elastin.

### 6. Sircol collagen assay

We assessed collagen content of the collagen using Sircol Assay Kits (Biocolor Ltd., Carrickfergus, UK). Actual procedures were according to the instructions of the manufacturer. Cells in 96-well plates with serum-free DMEM were co-cultured with various concentration of Feminilove vaginal gels. Cell lysates and supernatants were collected 48 hours after the initiation of experiment.

### 7. Statistical analysis

Statistical analyses were performed using GraphPad Prism version 9.1.1. All data are presented as mean ± S.E.M. For comparison of the means between two groups, data were analyzed by Student’s 2-tailed t test. Differences of the means for more than two groups containing two variables were analyzed using 2-way ANOVA. *Post-hoc* analysis was performed with Tukey’s test. A *p* value less than 0.05 was considered significant.

## III. RESULTS

### 1. Minimal toxicity of vaginal gel products

Detailed information for 4 vaginal gel products (Vmagic Organic Vulva Cream vaginal moisturizer, Vulvacare vaginal moisturizer & intimate skin cream, Membrasin Topical Vulua Cream for Feminine Dryness and Feminilove BIO-FRESH moisturizing vaginal gel) are listed in Table 1.

Our results suggest that all four vaginal gel products exhibited a minimal toxicity on cultured human vaginal cells (Figure 1). More importantly, toxicity tolerance of Feminilove vaginal gel is similarly to those 3 most popular vaginal gel products available in Amazon (Vmagic Organic Vulva Cream vaginal moisturizer, Vulvacare vaginal moisturizer & intimate skin cream and Membrasin Topical Vulua Cream for Feminine Dryness).

**Figure 1:**
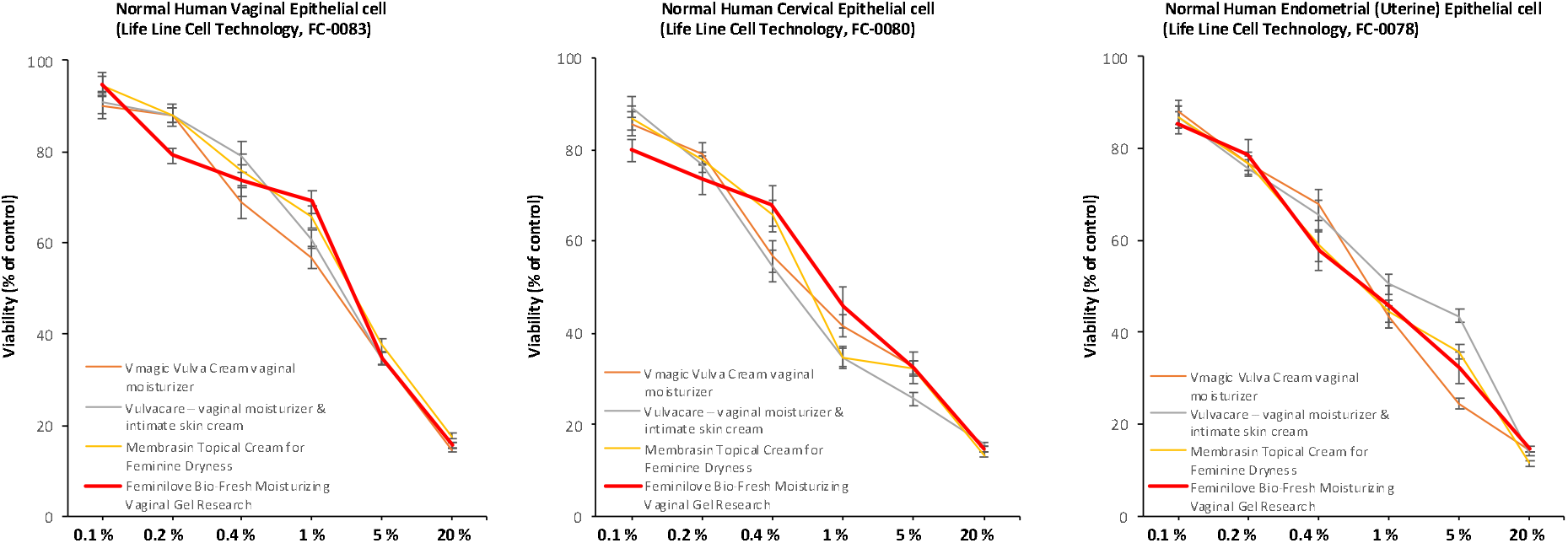
Cellular viability profiles for various products. Results are presented as the means and bars represent standard deviation from 3 independent experiments in which each condition was tested in triplicate.

### 2. Effect of Feminilove vaginal gel on the rabbit’s vaginal mucosa

Table 2 listed the results of histological morphology in the rabbit vaginal tissue after 10 days of intravaginal application of a gel formulation with or without 0.01, 0.1, or 1% Feminilove vaginal gel.

**Table 2:** Score of vaginal irritation for NZW rabbit.

|  |  | <u>Feminilove vaginal gel</u> |  |  |
| --- | --- | --- | --- | --- |
| Vaginal component (range of possible score) | Gel control (n=6) | 0.01% (n=6) | 0.1% (n=6) | 1% (n=6) |
| Epithelial exfoliation | 0.13 ± 0.04 | 0.33 ± 0.27 | 0.94 ± 0.41 | 0.33 ± 0.54 |
| Vascular congestion | 0.71 ± 0.22 | 0.76 ± 0.12 | 0.77 ± 0.17 | 1.75 ± 0.30 |
| Leukocyte infiltration | 1.02 ± 0.12 | 1.43 ± 0.32 | 1.05 ± 0.15 | 1.25 ± 0.13 |
| Lamina propra thickness | 0.95 ± 0.14 | 0.76 ± 0.24 | 1.05 ± 0.20 | 0.91 ± 0.33 |
| Total score | 2.81 | 3.28 | 3.81 | 4.24 |

Figure 2 shows representative mid vaginal sections of rabbits repeatedly exposed to placebo control, 0.01%, 0.1% and 1.0% of Feminilove vaginal gel. Feminilove vaginal gel showed acceptable vaginal irritation (total score 1–6).

**Figure 2:**
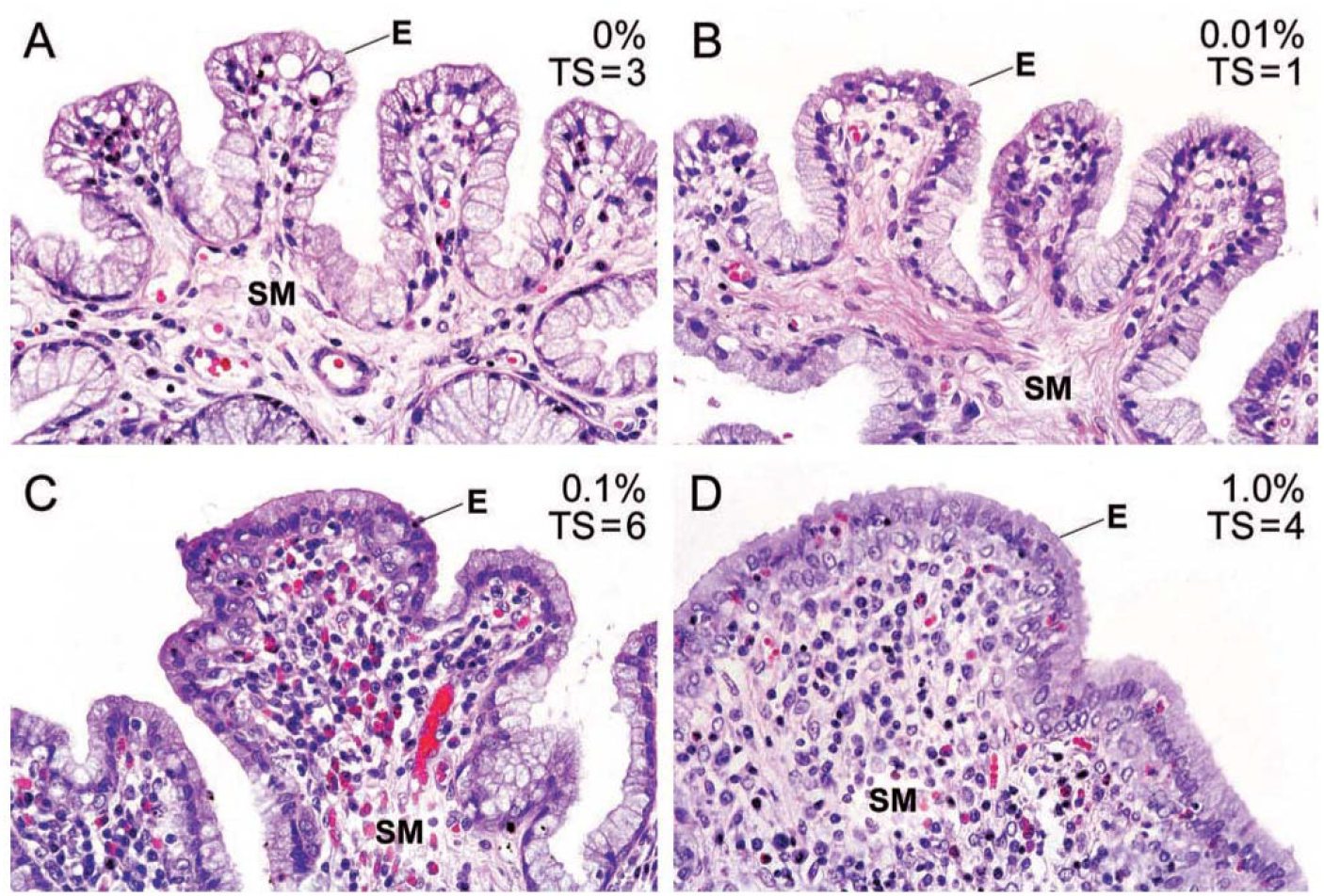
Light microscopy images of Feminilove vaginal gel-treatment rabbit vaginal sections. Original magnification × 400.

In addition, Table 3 lists the effects of Feminilove vaginal gel (0.01, 0.1, or 1.0%) on PCNA-positive cells in the vaginal epithelium and stromal cells of tissue sections from rabbits. We showed that > 24% and > 16% of epithelial cell and stromal cells, respectively, were stained for positive PCNA cells. Importantly, there was no significant difference in PCNA values between placebo control treated versus Feminilove vaginal gel treated samples.

**Table 3:** Cell proliferation results Feminilove gel for 10 days. The percentage of PCNS-positive cells were determined by light microscopy.

| Treatment | PCNA-positive cells (%) |  |
| --- | --- | --- |
|  | Epithelium | Stroma |
| Placebo group (n=6) | 24.5 ± 10.6 | 16.8 ± 5.6 |
| Feminilove treatment |  |  |
| 0.01% (n=6) | 34.9 ± 2.6 | 21.7 ± 4.9 |
| 0.1% (n=6) | 33.3 ± 3.5 | 26.5 ± 4.5 |
| 1% (n=6) | 24.6 ± 4.5 | 23.6 ± 4.8 |

### 3. Minimal systemic toxicity of Feminilove vaginal gel

We evaluated hematology parameters in rabbits treated with Feminilove vaginal gels versus placebo controls (Table 4). Importantly, hematology parameters between these 2 groups of rabbits were not different.

**Table 4:** Hematological results for rabbits treated with Feminilove vaginal gel application versus placebo control.

| Parameter | Gel control | Feminilove, 1% |
| --- | --- | --- |
| RBC x 10<sup>4</sup> / ul | 575 ± 25 | 587 ± 29 |
| WBC x 10<sup>3</sup> / ul | 3.2 ± 0.6 | 3.7 ± 0.9 |
| LYM % | 50.1 ± 11.4 | 45.8 ± 10.5 |
| NEU % | 42.7 ± 10.6 | 45.7 ± 8.9 |
| Others % | 7.1 ± 3.3 | 6.9 ± 3.7 |
| HGB g/dl | 11.4 ± 0.6 | 12.1 ± 0.8 |
| HCT % | 37.0 ± 2.7 | 42.3 ± 2.1 |
| MCV fl | 64.3 ± 4.5 | 65.9 ± 4.7 |
| MCH pg | 19.9 ± 0.6 | 21.5 ± 2.1 |
| MCHC g/dl | 41.6 ± 1.4 | 36.9 ± 3.8 |
| RDW % | 15.7 ± 1.4 | 15.8 ± 2.6 |
| RLT x 10<sup>4</sup> / ul | 897 ± 165 | 983 ± 146 |
| PMPV fl | 5.2 ± 0.7 | 5.6 ± 2.2 |

### 4. Minimal cell toxicity of Feminilove vaginal gel on reconstituted human vaginal epithelium model

In 3 models of reconstituted human vaginal cells (epithelial, cervical and endocervical epithelial cells), co-cultured with various concentration of Feminilove vaginal gel (0.01, 0.1, and 1.0%) for 24 hours did not show sign of cytotoxicity (Figure 3, left panels). Result of cell culture study is in agreement with the data when topically Feminilove vaginal gel has been applied for 10 days in female rabbits (Figure 3, right panels). Both *in vitro* cell culture study and *in vivo* rabbit study demonstrated the minimal cell toxicity of Feminilove vaginal gel.

**Figure 3:**
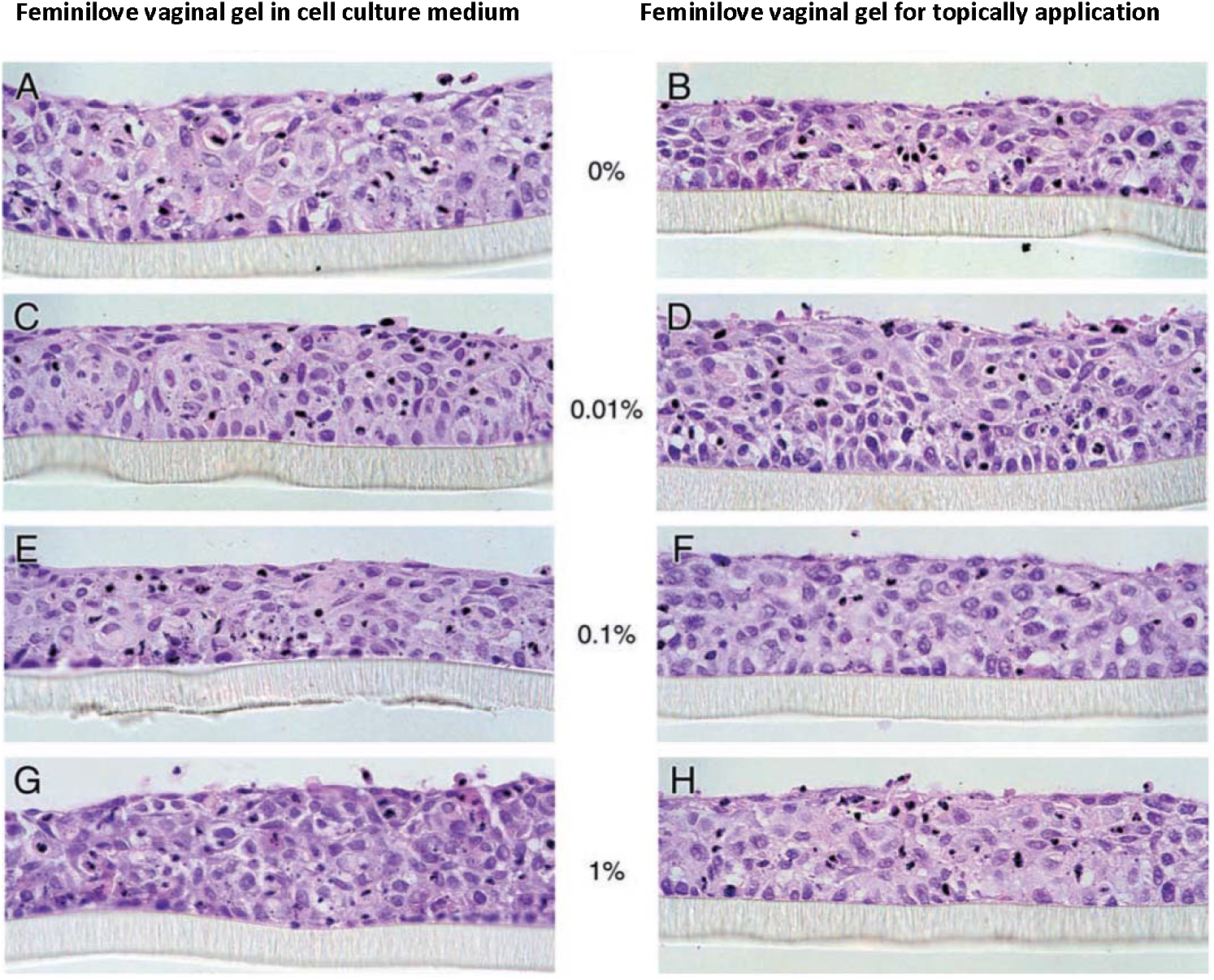
Lack of membrane damaging effect of Feminilove vaginal gel on reconstituted human vaginal epithelia. in vitro 3-D tissue cultures of human vaginal epithelial cells grown in polycarbonate membrane filters were exposed to Feminilove vaginal gel in culture medium (left column) or topically via gel (right column) and toxicity was evaluated by histological assessment of H&E-stained tissue sections. Cultures were exposed to 0, 0.01, 0.1, or 1% Feminilove vaginal gel for 24 hours.

### 5. Regulatory effects of Feminilove vaginal gel on the growth of uropathogenic and commensal vaginal bacteria

A significant large proportion of post-menopausal women have applied vaginal lubricating products to relieve post-menopause associated genitourinary syndrome. We compared the effects of 3 common vaginal products (Vmagic Organic Vulva Cream vaginal moisturizer, Vulvacare vaginal moisturizer & intimate skin cream, Membrasin Topical Vulua Cream for Feminine Dryness) versus Feminilove vaginal gel on growth and viability of cultured *Escherichia coli* and *Lactobacillus crispatus*. Feminilove vaginal gel significantly inhibited growth of *Escherichia coli* (Figure 4A). Vaginal *Lactobacillus* species inhibit growth of *Escherichia coli*. We showed that among 4 vaginal gel products, only Feminilove vaginal gel promoted the growth of *Lactobacillus spp*. (Figure 4B).

**Figure 4:**
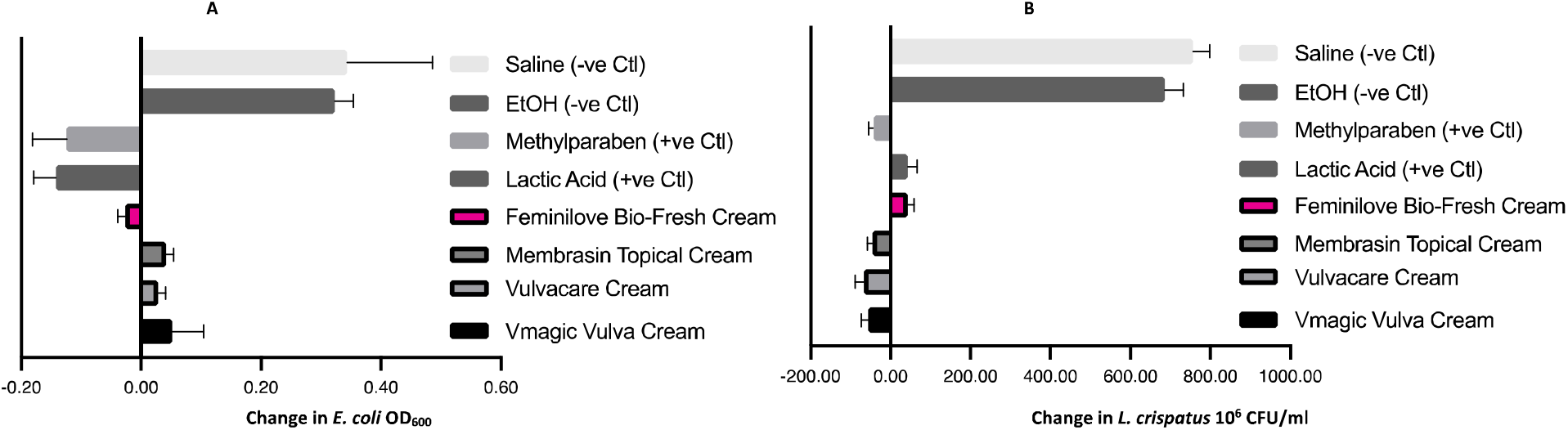
Change in bacterial growth with exposure to commercial vaginal moisture products. (A) Final change in growth as measure by OD600, for laboratory strain of E.coli. All products exhibit similar inhibitory properties of E. coli. (B) Final change in growth of L. crispatus, as measured in CFU (simple co-culture without normal human vaginal epithelial cell, Line Cell Technology, FC-0083). Results are presented as the means and bars represent standard deviation from 3 independent experiments in which each condition was tested in triplicate.

### 6. Anti-inflammatory effects of Feminilove vaginal gel in cultured human vaginal epithelial cells

Inflammation is often present in the female reproductive tract. We showed that Feminilove vaginal gel elicits an anti-inflammatory response and significantly decreased inflammatory mediators (IL-6, TNF-α and IL-1β) induced by LPS on cultured vaginal epithelial cells (Figure 5).

**Figure 5:**
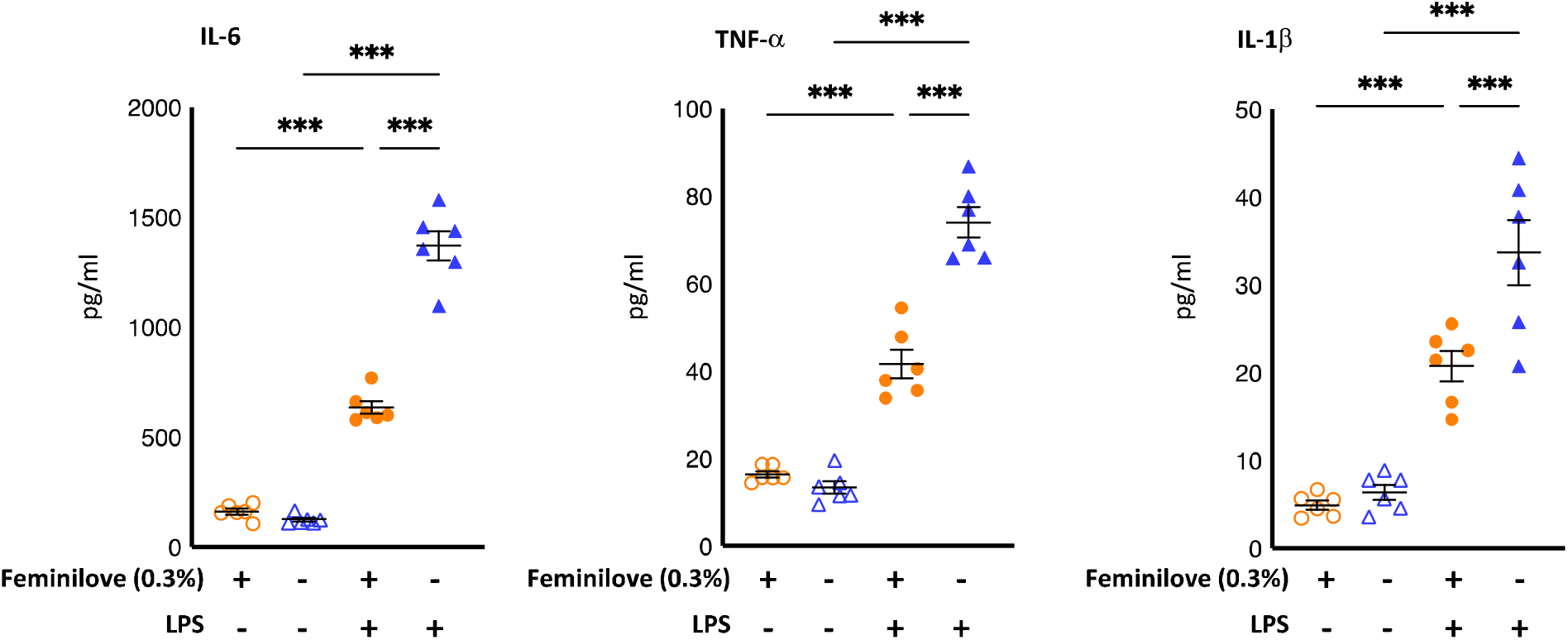
Feminilove vaginal gel elicits an anti-inflammatory response and decreases inflammatory mediators induced by the stimulation of lipopolysaccharides (LPS) for 18 hours at pH 3.9 as indicated. The presence of IL-6, TNF- and IL-1 was measured in the supernatant collected at the end of the study. Concentration of inflammatory cytokines were determined by a Luminex-based cytokine assay. *** denotes p < 0.001.

### 7. Production of surface-deposited tropoelastin and collagen in cultured vaginal smooth muscle cells

The vaginal wall and structural support of vaginal pelvis is related to content of collagen, elastin, and muscle. We showed that Feminilove vaginal gel significantly increased tropoelastin and collagen production in cultured vaginal smooth muscle cells (Figure 6).

**Figure 6:**
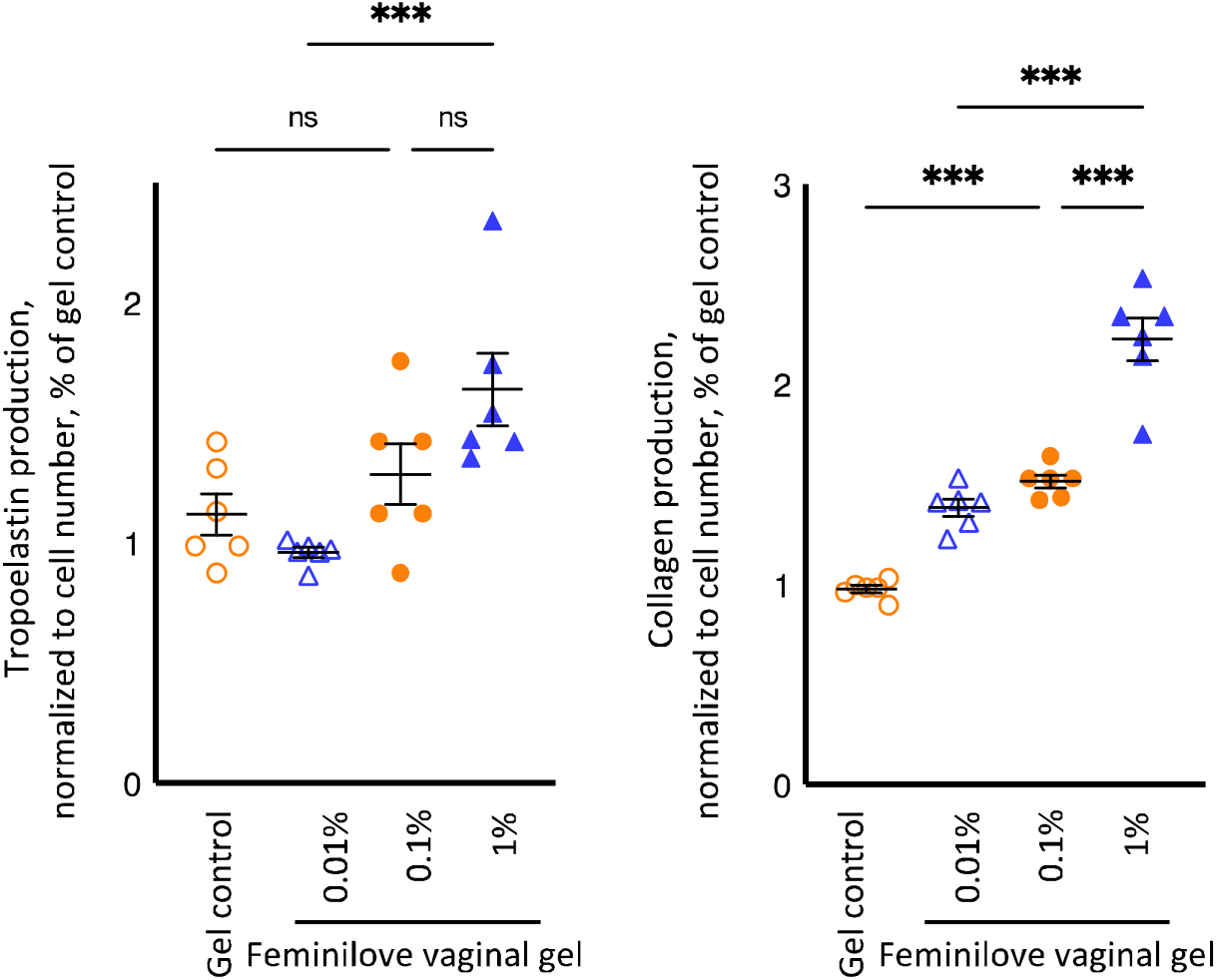
Effects of Feminilove vaginal gel on tropoelstin and collagen (cell culture media) production on vaginal smooth muscle cells at 48 hours. Significant increase of tropoelatin and collagen production was observed at 1% and 0.1% of Feminilove vaginal gel, respectively.

## IV. DISCUSSION

Vaginal dryness is common in women and has a significant negative impact on their sexual satisfaction and overall quality of life. Due to cultural influence, this condition is often underreported.^26^ In this study, we evaluated the safety and the beneficial effects of Feminilove BIO-FRESH moisturizing vaginal gel, a new type of estrogen-free vaginal gel. Our results suggest that; 1) Feminilove vaginal gel exhibits minimal cell cytotoxicity on various human vaginal cells; 2) Feminilove vaginal gel exhibits minimal side-effects on the structure of vaginal mucosa stratum of experimental animals; 3) Feminilove vaginal gel inhibits the growth of pathogenic vaginal bacteria (*E. coli*) while promotes the growth of beneficial vaginal bacteria (*Lactobacillus spp*); 4) Feminilove vaginal gel demonstrates an anti-inflammatory response on cultured vaginal epithelial cells; and 5) Feminilove vaginal gel promotes the production of tropoelastin and collagen on cultural vaginal smooth muscle and thus, may restore loose vaginal wall (*i.e.*, tightening effects).

One of the important ingredients of Feminilove vaginal gel is aloe vera. Aloe vera is a natural product that have been frequently used in the field of cosmetology. The aloe vera plant has been used for centuries for its beneficial effects. Aloe vera contains numerous potentially active constituents such as vitamins, enzymes, mineral, sugar, lignin, saponins, salicylic acids and amino acids. The mechanism of its beneficial action includes healing properties, effects on skin rejuvenation, anti-inflammation, modulatory effects on the immune system, anti-tumor, as well as moistening and anti-aging activities.^27^ In addition, aloe vera exerts inhibitory action on fungi, bacteria, and viruses. Moreover, aloe vera gel possesses moisturizing and anti-aging effects. Mucopolysaccharides from aloe vera extract promotes the binding of moisture into the skin. Aloe promotes fibroblast to produce collagen and elastin and as a result, exerts its anti-aging and anti-wrinkle effects.^28,29^

## V. CONCLUSIONS

The negative impact of vaginal dryness on quality of life is overwhelming. In our *in vitro* and *in vivo* models, we showed that Feminilove BIO-FRESH moisturizing vaginal gel is a safe and effective remedy for women with vaginal dryness and vulvovaginal atrophy.

## Acknowledgements

Financial support for this study was provided by the Epoch NE Corporation, Seattle, WA. However, the funder has no role in the design of the study; in the collection, analyses, or interpretation of the data; in the writing of the manuscript, or in the decision to publish the results. Design of this work has followed the study guidelines from the American Gynecological & Obstetrical Society. The American Gynecological & Obstetrical Society is an organization composed of individuals attaining national prominence in scholarship in the discipline of Obstetrics, Gynecology and Women’s Health as well as to advance the health of women by providing dedicated leadership and promoting excellence in research, education and medical practice.

## References

1. Dunn KM, Croft PR, Hackett GI. Sexual problems: a study of the prevalence and need for health care in the general population. Fam Pract 1998;15:519–24

2. Leiblum SR, Hayes RD, Wanser RA, Nelson JS. Vaginal dryness: a comparison of prevalence and interventions in 11 countries. J Sex Med 2009;6:2425–33

3. Palacios S. Managing urogenital atrophy. Maturitas 2009;63:315–8

4. Panay N, Fenton A. Vulvovaginal atrophy – a tale of neglect. Climacteric 2014;17:1–2

5. Portman DJ, Gass ML, Vulvovaginal Atrophy Terminology Consensus Conference Panel. Genitourinary syndrome of menopause: new terminology for vulvovaginal atrophy from the International Society for the Study of Women’s Sexual Health and The North American Menopause Society. Climacteric 2014;17:557–63

6. Panay N. Genitourinary syndrome of the menopause – dawn of a new era. Climacteric 2015;18(Suppl 1):13–7

7. Braunstein S, van de Wijgert J. Preferences and practices related to vaginal lubrication: implications for microbicide acceptability and clinical testing. J Womens Health (Larchmt) 2005;14:424–33

8. Jozkowski KN, Herbenick D, Schick V, Reece M, Sanders SA, Fortenberry JD. Women’s perceptions about lubricant use and vaginal wetness during sexual activities. J Sex Med 2013;10:484–92

9. Andelloux M. Products for sexual lubrication: understanding and addressing options with your patients. Nurs Womens Health 2011;15:253–7

10. Cordeau D, Courtois F. Sexual disorders in women with MS: assessment and management. Ann Phys Rehabil Med 2014;57:337–47

11. Kennedy SH, Rizvi S. Sexual dysfunction, depression, and the impact of antidepressants. J Clin Psychopharmacol 2009;29:157–64

12. Sutton KS, Boyer SC, Goldfinger C, Ezer P, Pukall CF. To lube or not to lube: experiences and perceptions of lubricant use in women with and without dyspareunia. J Sex Med 2012;9:240–50

13. Jamieson DJ, Steege JF. The prevalence of dysmenorrhea, dyspareunia, pelvic pain, and irritable bowel syndrome in primary care practices. Obstet Gynecol 1996;87:55–8

14. Seehusen DA, Baird DC, Bode DV. Dyspareunia in women. Am Fam Physician 2014;90:465–70

15. Lev-Sagie A. Vulvar and vaginal atrophy: physiology, clinical presentation, and treatment considerations. Clin Obstet Gynecol 2015;58:476–91

16. Kingsberg SA, Wysocki S, Magnus L, Krychman ML. Vulvar and vaginal atrophy in postmenopausal women: findings from the REVIVE (REal Women’s View’s of Treatment Options for Menopausal Vaginal ChangEs) survey. J Sex Med 2013;10:1790–9

17. Goldstein AT, Burrows LJ. Sexual Pain Disorders in Women 2009 [7 July 2015]. Available from: http://www.issm.info/news/review-reports/sexual-pain-disorders-in-women/

18. Sturdee DW, Panay N. Recommendations for the management of postmenopausal vaginal atrophy. Climacteric 2010;13:509–22

19. The North American Menopause Society. Management of symptomatic vulvovaginal atrophy: 2013 position statement of The North American Menopause Society. Menopause 2013;20:888–902

20. Chen J, Geng L, Song X, Li H, Giordan N, Liao Q. Evaluation of the efficacy and safety of hyaluronic acid vaginal gel to ease vaginal dryness: a multicenter, randomized, controlled, open-label, parallel-group, clinical trial. J Sex Med 2013;10:1575–84

21. Bachmann GA, Nevadunsky NS. Diagnosis and treatment of atrophic vaginitis. Am Fam Physician 2000;61:3090–6

22. Sinha A, Ewies AA. Non-hormonal topical treatment of vulvovaginal atrophy: an up-to-date overview. Climacteric 2013;16:305–12

23. https://www.promega.com/-/media/files/resources/protocols/technical-manuals/101/celltiterglo-2-0-assay-protocol.pdf?la=en

24. Eckstein, P., Jackson, M. C., Millman, N., and Sobrero, A. J. Comparison of vaginal tolerance tests of spermicidal preparations in rabbits and monkeys. J Reprod Fertil 1969;20:85–93

25. D’Cruz, O. J., and Uckun, F. M. Intravaginal toxicity studies of a gelmicroemulsion formulation of spermicidal vanadocenes in rabbits. Toxicol Appl Pharmacol 2001;170:104–12

26. Willhite LA, O’Connell MB. Urogenital atrophy: prevention and treatment. Pharmacotherapy 2001;21:464–80

27. Surjushe A, Vasani R, Saple DG. Aloe vera: a short review. Indian J Dermatol 2008;53:163–6

28. Shelton M. Aloe vera, its chemical and therapeutic properties. Int J Dermatol 1991;30:679–783

29. West DP, Zhu YF. Evaluation of aloe vera gel gloves in the treatment of dry skin associated with occupational exposure. Am J Infect Control 2003;31:40–2

